# The forest avifauna of Arabuko Sokoke Forest and adjacent modified habitats

**DOI:** 10.1101/2021.04.19.440426

**Authors:** David O. Chiawo, Wellington N. Kombe, Adrian J.F.K. Craig

## Abstract

Arabuko Sokoke Forest (ASF) is the largest area of coastal forest remaining in East Africa and a major Important Bird Area in mainland Kenya. The study analysed data from point count surveys over 15 months in three land use types; primary forest (PF), plantation forest (PL), and farmlands (FM), and compared these to the first comprehensive bird checklist for the forest, as well as recent surveys from other studies. Avifaunal diversity and abundance were compared using multivariate analysis to determine bird responses to different land use characteristics. The primary forest held a distinctive bird community, while the bird communities of farmlands and plantation forest were more similar to each other. Land use had a significant effect on overall avian diversity and abundance. The current forest avifauna was divided into forest specialists (16 species), forest generalists (26 species) and forest visitors (30 species). Seven species of forest specialist and generalists recorded prior to 1980 may no longer occur in the forest. Of 38 specialists and generalists recorded in our point counts, 19 were also recorded on farmland and 28 in plantations. One forest specialist, the Green Barbet, was most encountered outside the forest. Future research should focus on habitat use by these bird species, and the extent of movement by forest birds between the remaining patches of coastal forest. Patterns of habitat use by birds in the area suggest that vegetation heterogeneity and habitat complexity are especially significant in sustaining diverse and abundant bird populations. The management of plantations and farmland will be critical for the conservation of forest generalists and forest visitors.

## INTRODUCTION

Since protected areas alone will be inadequate to conserve global biodiversity including birds, the management of human-modified habitats is critical for tropical bird conservation (1). Predicting the response of bird communities to habitat disturbances is still a challenge (2). Past studies have indicated a strong relationship between forest area and the number of species of forest birds (3). While agroforests with a mix of cultivated and natural shade trees will attract a considerable number of bird species(1,4), there may be a marked reduction in forest visitors and overall shifts in species composition have been reported in agroforestry systems (5,6). As habitats become more disturbed and open, forest-dependent species are replaced by generalists and open-area species (5,6) with a marked shift in assemblage composition (6). Thus even small changes in the structure and composition of the tree cover may have a significant impact on bird assemblages (5), leading to changes in bird diversity and community composition (7). Habitat changes particularly affect less abundant and range-restricted birds, rainforest specialists and altitudinal migrants (8). Apart from habitat specialization, life history traits such as large territories and large body size (9) may also influence species vulnerability. Arabuko Sokoke Forest (ASF), with a total area of approximately 41 600 ha, is the largest surviving single block of the previously extensive indigenous coastal tropical forest in Eastern Africa. It is situated in Coastal Kenya at, 3° 20’ S and 39° 50’ E, between Kilifi and Malindi, 110 km north of Mombasa. It was proclaimed a Crown Forest in 1932, and gazetted as a forest reserve in 1943 during the colonial period, then designated a strict nature reserve in the 1960s (10). Arabuko Sokoke Forest is now surrounded on all sides by 54 village communities with a total population of about 130 000 people (11). The reduction of ASF and modification of adjacent habitats can be traced to beyond 1960s due to licensed and unlicensed cutting of timber trees (12). However, the levels of unsustainable forest use have intensified, with increasing human populations resulting in higher levels of dependence on the forest for both domestic and commercial resources (13). Meanwhile, Arabuko Sokoke Forest is rapidly gaining a reputation in ecotourism for tourists with an interest in birds. The concentration of rare bird species accounts for its status as the second most important forest for the conservation of threatened bird species on the African mainland (10). A total of 261 bird species has been recorded within the forest (14) with six globally threatened species: Clarke’s Weaver (*Ploceus golandi*), Sokoke Scops Owl (*Otus ireneae*), Amani Sunbird (*Anthreptes pallidigaster*), East Coast Akalat (*Sheppardia gunningi*), Spotted Ground Thrush (*Zoothera guttata*) and Sokoke Pipit (*Anthus sokokensis*). It also protects other taxa, such as distinctive near-endemic small mammals, and many endemic invertebrates.

An earlier study examined changes in the avifauna of the forest and adjoining habitats, related to land use at the level of bird feeding guilds (15). In this paper, we focus on the species level and examine the effects of land use on the composition of the bird community, in three land use types; primary forest (PF), plantation forest (PL), and farmlands (FM). The primary forest consisted of three habitat types including *Brachystegia*, Mixed Forest and *Cynometra*. The responses of different species to habitat modification can inform land use management for bird conservation in the area, and the maintenance of this forest as an Important Bird Area (IBA) and a potential Key Biodiversity Area (KBA).

## MATERIALS AND METHODS

### Data collection

The surveys covered three land use types: primary forest, plantation forest, and farmlands. Farmlands were characterized by smallholder farming of maize and cassava; bush clearing, burning, logging of trees, and few tree stands. Primary forest had a high vegetation heterogeneity due to the three habitat types: *Brachystegia* and Mixed Forest had a higher vertical heterogeneity due to many large trees and a dense understory, compared to *Cynometra* which had no large trees, but thicket at a uniform height. Plantation forest consisted of trees for commercial timber including *Eucalyptus* sp. and *Casuarina* sp., with open areas due to timber harvesting. *Eucalyptus* sp. stands were 8-10 m high and were mature for harvesting, while *Casuarina* were young stands.

We used point counts as described by (16) to survey birds. Calling distance was estimated from the radial distance from the point to the bird (17). The coordinates of point count stations on each route were recorded using a Garmin 20 GPS. Three transects per land use type were set up according to habitat heterogeneity in farmland and plantation forest, and by vegetation type in the primary forest. Points were spaced 100 m apart along each transect in an alternating manner on either side of each transect. The points were located 30m off the track in the primary forest to minimize edge effects, due to the track through the forest. There were no replication plots in this study apart from the three transects in every land use type.

Bird counts lasted for 20 minutes at each point in all the three land use types. We used a longer time (additional 5 minutes) rather than the commonly-used 15 minutes to increase the likelihood of recording inconspicuous species in the dense vegetation as suggested by (18). All birds seen or heard within a 50 m radius were recorded; their distances from the centre of the point count station and the perching heights were estimated. The radius was selected based on the Effective Detection Radius (EDR) that has been used in many surveys comparing forest and farmland birds in Kenya

e.g. (19,20). Birds seen or heard beyond this radius were ignored and did not form part of our data. Birds detected only in flight were also excluded from the data and did not form part of the analysis. Surveys were on days without persistent or heavy rain from 0630 - 1100 when birds were most active.

The direction of travel through counts was rotated to minimize any potential bias from the time of day. Transects followed established footpaths and forest tracks and where possible at least nine points were located along each one km transect to standardize survey effort. In total 81 point count stations were surveyed along nine transects, three in each land use type; farmland (FM), Primary forest (PF), and Plantation (PL). Each point count station was surveyed once on each monthly visit during the whole survey period from May 2012 to September 2013, covering dry and wet seasons.

Points for each land use type were surveyed on alternate days; therefore, points for each land use category were randomized separately. The process was repeated every evening before the next day of the survey until all points were surveyed in every month of the field visits. This limited movement to short distances between points in one land use type on each morning of the survey and avoided long travelling time between points. Points within the primary forest were distributed equally in the three vegetation types (Mixed Forest, *Brachystegia* woodland and *Cynometra*), nine points in each vegetation type. An equal number of points (n=27) from forest, farmland and plantation forest ensured standard sampling effort for comparison. A standard area (‘covered area’) of 0.21 km^2^ was sampled in each land use type. Therefore, the areas sampled were constant in the three land use types to allow comparison of species richness and diversity. Land use type was the unit of analysis upon which the comparisons were made. The area surveyed was 4.8 km^2^ in primary forest, 5.4 km^2^ in plantation forest and 6.4 km^2^ in farmland, which we used in calculating bird abundance. The ‘area covered’ was the total area covered by the 27 points within the area surveyed. While the “area covered” was standard for all the three habitats, their dimensions were different, hence, differences in the total area surveyed.

Vertical perching height of birds detected by both calls and sight was estimated within the limits of 0-3 m, 3-12 m, and > 12 m. The total number of individual birds seen and heard at each point count station was considered to be the count of individuals at each visit. Individual counts from each point were pooled per land use type to obtain the total counts of each species. To obtain data on habitat structure, we recorded vertical vegetation heterogeneity, the number of large trees, the proximity of each point to settlements and to the forest, and the number of trees with fruits. The number of fruiting trees were counted within a standard area of 50 m radius circle at each point. To determine vertical vegetation heterogeneity at each point, plant cover was estimated to the nearest 5% at heights of 0, 1, 2, 4, 8 and 16 m (21). Vertical vegetation heterogeneity was then defined as the diversity of vegetation layers using the Shannon–Wiener diversity index (16,21). Birds recorded were classified into three categories according to forest utilization: forest specialists (FF), forest generalists (F) and forest visitors (f) according to (22).

### Data analysis

Species richness was calculated as the cumulative number of species recorded at each point count station. ANOVA and Tukey HSD post hoc tests were used to test for differences in bird diversity and abundance among the land use types. Species diversity at each point was calculated based on the Shannon diversity index (*H*) (23)

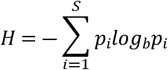

We used General Linear Mixed Models (GLMM) to test the mixed effect of habitat factors on richness of birds based on Akaike’s Information Criterion (AIC), the model with the lowest AIC value is considered to be the best (21,24). This approach is considered a useful tool for model selection in ecology (21). All statistical analysis was done with R statistical software (25).

We estimated abundance of each bird species encountered in each land use by multiplying bird density by the area of each land use surveyed (26);

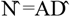

Where, 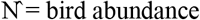, A= total area of land use surveyed and 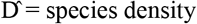.

Species density 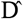 was estimated by:

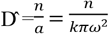

Where k=total number of points in each land use surveyed, n= number of birds (for each species of interest) counted summed across all points, ω = the fixed radius of the point, and a = *kπ*ω^2^= the total area of the surveyed points, termed the ‘covered area’. We performed non-metric multidimensional scaling to establish similarity in bird community composition between sampling points.

## RESULTS

### Species composition

Over the whole survey period, a total of 97 bird species was recorded at 81 points (Appendix 1). A Jacknife procedure estimated the expected bird species richness for the area covered by the counts to be 114.82. Thus, the number of species obtained was close to the expected, suggesting that the counts had sampled the whole bird community.

Of the 29 species in the East African Coastal forest which occur in Kenya, 26 have been recorded in Arabuko Sokoke (10,14). During the point counts 17 of these species were recorded (Table 1). Eight forest specialists (FF) were recorded mainly in the primary forest: Fischer’s Greenbul (*Phyllastrephus fischeri*), Tiny Greenbul (*Phyllastrephus debilis*), Sokoke Pipit (*Anthus sokokensis*), Pale Batis (*Batis soror*), East Coast Akalat (*Sheppardia gunningi*), Four-coloured Bush-shrike (*Malaconotus quadricolor*), Amani Sunbird (*Anthreptes pallidigaster*) and Clarke’s Weaver (*Ploceus golandi*). Three of these species, Clarke’s Weaver, Sokoke Pipit, and East Coast Akalat, are Red Listed as globally threatened (27). Thirteen species were never recorded on farmland, and only one, the Scaly Babbler (*Turdoides squamulatus*), was common in this habitat and in fact not recorded in the primary forest during point counts. Six of the seventeen species were not recorded from the plantation areas, but some appeared to use this habitat regularly for foraging: Southern Banded Snake Eagle (*Circaetus fasciolatus*), Mangrove Kingfisher (*Halcyon senegaloides*), Eastern Green Tinkerbird (*Pogoniulus simplex*), Mombasa Woodpecker (*Campethera mombassica*), Scaly Babbler, Little Yellow Flycatcher (*Erythrocercus holochlorus*), Chestnut-fronted Helmet-shrike (*Prionops scopifrons*), and Black-bellied Starling (*Notopholia corusca*).

**Table 1:** Birds restricted to the East African Coastal forest recorded during point counts. Their observed abundance in different habitats (cf. Appendix 1) is indicated by 1 (< 10 records) 2 (10-20 records) 3 (21-50 records), 4 (> 50 records). FF=Forest specialists, F=Forest generalists, f=forest visitors. Categorization follows the classification of forest birds by (22).

| Category | Species | Forest | Plantation | Farmland |
| --- | --- | --- | --- | --- |
| FF | Fischer's Greenbul *Phyllastrephus fischeri* | 4 | 1 | 0 |
| F | Pale Batis *Batis soror* | 3 | 0 | 0 |
| F | Chestnut-fronted Helmet-shrike *Prionops scopifrons* | 3 | 2 | 0 |
| F | Four-coloured Bush-shrike *Malaconotus quadricolor* | 3 | 0 | 0 |
| FF | Tiny Greenbul *Phyllastrephus debilis* | 3 | 0 | 0 |
| FF | Little Yellow Flycatcher *Erythrocercus holochlorus* | 3 | 3 | 0 |
| FF | Black-bellied Starling *Notopholia corusca* | 3 | 3 | 2 |
| FF | Amani Sunbird *Anthreptes pallidigaster* | 3 | 1 | 0 |
| F | Mombasa Woodpecker *Campethera mombassica* | 2 | 2 | 1 |
| F | Plain-backed Sunbird *Anthreptes reichenowi* | 2 | 1 | 0 |
| FF | Sokoke Pipit *Anthus sokokensis* | 2 | 0 | 0 |
| FF | East Coast Akalat *Sheppardia gunningi* | 2 | 0 | 0 |
|  | Mangrove Kingfisher *Halcyon senegaloides* | 1 | 1 | 1 |
| F | Southern Banded Snake Eagle *Circaetus fasciolatus* | 1 | 1 | 0 |
| FF | Eastern Green Tinkerbird *Pogoniulus simplex* | 1 | 1 | 0 |
| FF | Clarke's Weaver *Ploceus golandi* | 1 | 0 | 0 |
|  | Scaly Babbler *Turdoides squamulatus* | 0 | 3 | 3 |

Other forest specialists (FF) recorded in the primary forest were Plain-backed Sunbird (*Anthreptes reichenowi*), Forest Batis (*Batis mixta*), Blue-mantled Crested Flycatcher (*Trochocercus cyanomelas*), and Black-headed Apalis (*Apalis melanocephala*). Some forest specialists were recorded across the three habitat types including Green Barbet (*Stactolaema olivacea*), Collared Sunbird (*Hedydipna collaris*), and Black-bellied Starling.

Widespread forest visitors (f) such as Common Bulbul (*Pycnonotus barbatus*), Bronze Mannikin (*Lonchura cucullata*), Yellow-fronted Canary (*Crithagra mozambica*) and Fork-tailed Drongo (*Dicrurus adsimilis*) were common in platation forest and farmlands.

### Seasonal occurrence

The avifauna of ASF and surrounds includes few migrants (e.g. Black Kite (*Milvus migrans*), Lesser Striped Swallow (*Hirundo abyssinica*), something that this study confirms. (28) discussed evidence for altitudinal migration of birds from the Eastern Arc Mountains to coastal forests, based on their research in Tanzania. The Taita Hills, 200 km inland from Arabuko Sokoke Forest, represents the northernmost extension of the Eastern Arc Mountains, although they hold no species restricted to this Endemic Bird Area (10). (28) concluded that there was no evidence for migration by any of the endemic forest bird species in the Eastern Arc, but some indication of seasonal altitudinal movements for other montane forest birds. The only species on their list which has been recorded from ASF is the Eastern Bronze-naped Pigeon *Columba delegorguei* (14), but this bird was not encountered during our study. None of the bird species characteristic of the Afrotropical Highlands biome which have been recorded in the Taita Hills forests have been recorded in Arabuko Sokoke (10). Altitudinal migration has often been postulated for montane forest birds, but it may be a local phenomenon for many species (29). Ringing records from the Watamu area do indicate seasonal movements for the Red-capped Robin-Chat (*Cossypha natalensis*) within the coastal belt (Colin Jackson, pers. comm.).

### Species richness

Primary forest registered the highest bird mean species numbers (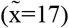) followed by plantation forest (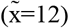), then farmlands (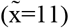). Contrary to our expectation plantation forest and farmlands had nearly equal species numbers (Fig. 2). We found a highly significant difference in species numbers between primary forest and farmland, Tukey test (*P*< 0.001), a significant difference between primary forest and farmland (*P*<0.05) but no significant difference between plantation and farmland (ns) (Fig 1).

**This is the Fig 1 legend:**
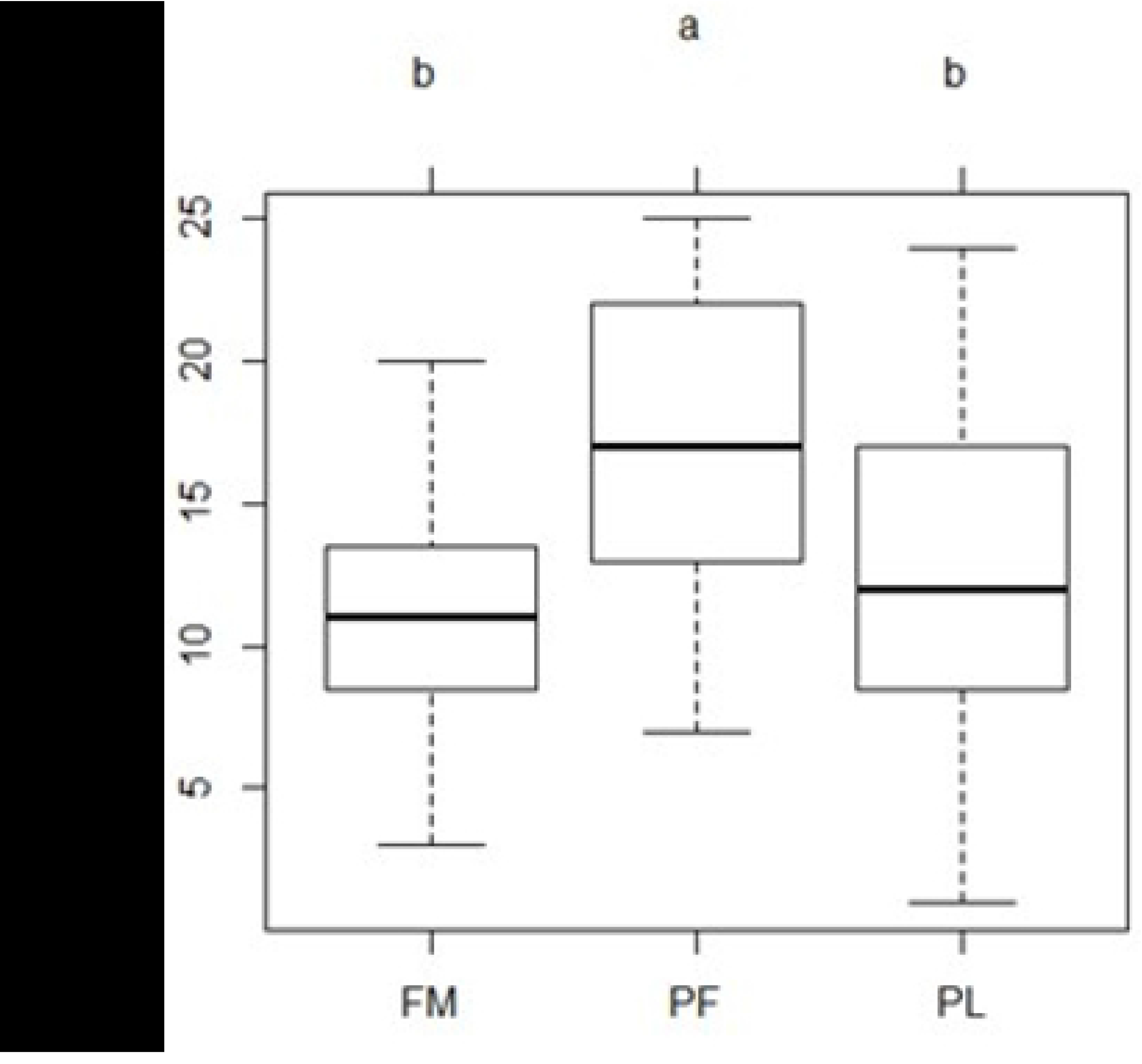
Box plots of mean avifauna richness per count station compared between primary forest (PF), farmland (FM), and Plantation forest (PL) at 95% CIs.

**This is the Fig 2 legend:**
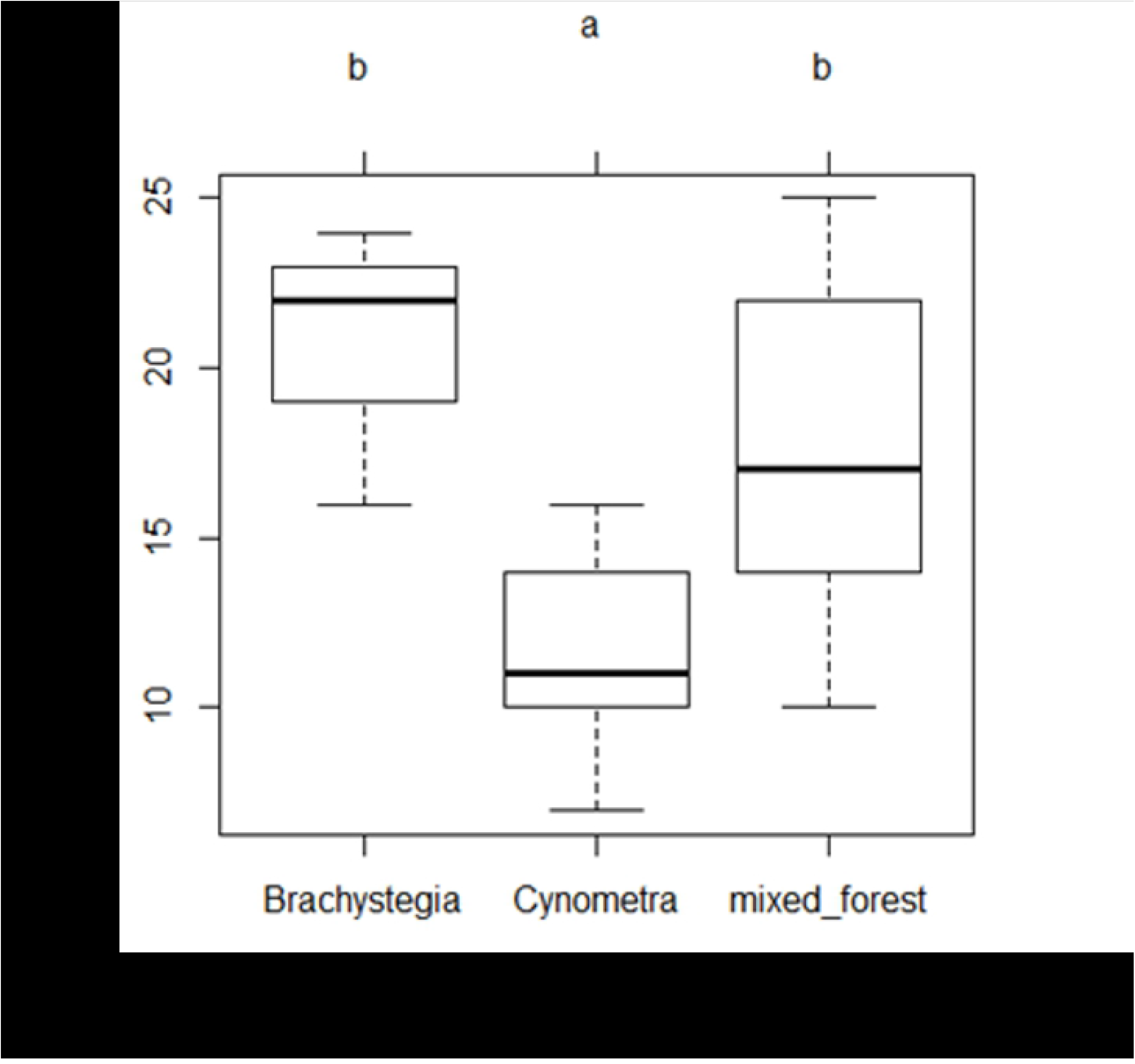
Box plots of mean bird species richness compared between three vegetation types (*Brachystegia, Cynometra* and Mixed Forest) in primary forest at 95% CIs.

Different vegetation types within the primary forest (*Brachystegia, Cynometra* and Mixed Forest) also showed significant differences in bird species composition, ANOVA (F2, 24 = 14.13, *P*< 0.001, n=27), with *Cynometra* vegetation having significantly lower species numbers than the other two. Multiple comparison of mean species composition using Tukey HSD revealed a highly significant difference between *Cynometra* and *Brachystegia* (*P*< 0.001) but no difference between Mixed Forest and *Brachystegia* woodland (ns) (Fig 2).

### Avian diversity and abundance

The highest species diversity (H) was recorded in primary forest (H = 2.75) and the lowest in farmland (H = 2.30). The plantation had an intermediate diversity (H = 2.48). Avian diversity was significantly different among the three land use categories, ANOVA (F_2, 78_ = 5.04, P = 0.009, n=81) (Fig 3). The Tukey test revealed significant differences between two pairs, primary forest and farmland and primary forest and plantation forest (*P*< 0.05).

**This is the Fig 3 legend:**
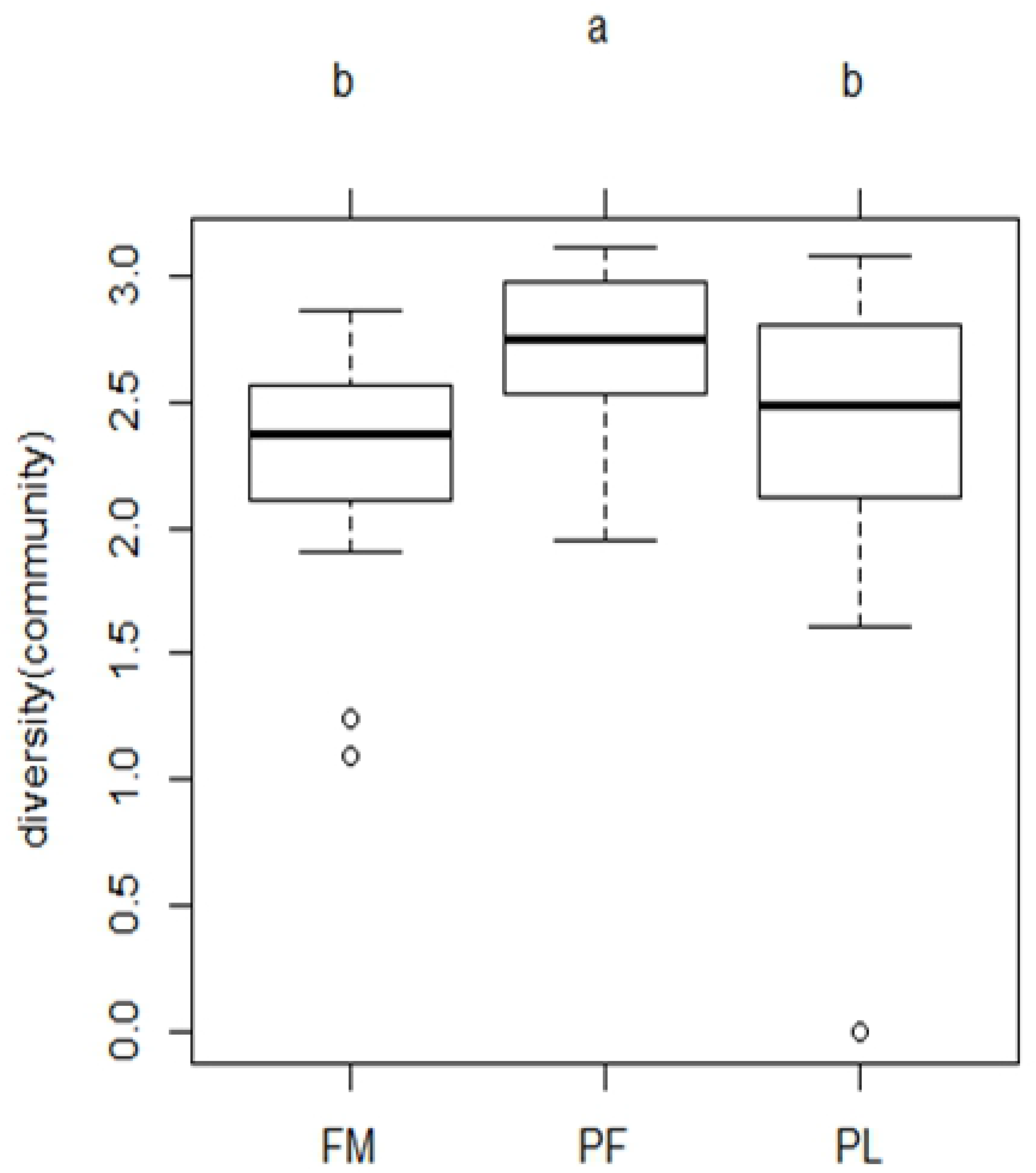
Box plots of mean Shannon diversity indices (H) in farmland (FM), primary forest (PF) and plantation forest (PL) at 95% CIs.

Some Forest specialists (FF) occurred in high calculated abundance (N>200) across the three land use types including Collared Sunbird, Olive Sunbird (*Nectarinia olivacea*), Black-bellied Starling, Green Barbet. The following species were most abundant in farmland and plantation forest including the Zanzibar Sombre Greenbul (*Andropadus importunus*), Yellow-fronted Canary, Bronze Mannikin, and Fork-tailed Drongo. Ashy Flycatcher (*Muscicapa caerulescens*), Northern Carmine Bee-eater (*Merops nubicus*), Grey-headed Kingfisher (*Halcyon leucocephala*), Green-backed Camaroptera (*Camaroptera brachyura*) and Red-billed Firefinch (*Lagonosticta senegala*) were recorded infrequently. Lesser Striped Swallow, Pallid Honeyguide (*Indicator meliphilus*), Lesser Honeyguide (*Indicator minor*), Scaly-throated Honeyguide (*Indicator variegatus*), Northern Brownbul (*Phyllastrephus strepitans*) and Retz’s Helmet-shrike (*Prionops retzi*) were most encountered in the plantation (Appendix I).

**Appendix I:** Classification and distribution of bird species by density and abundance in farmland, plantation and primary forest in Arabuko Sokoke Forest and neighboring land use types. n= total count of individual species, D^ =species density, N^ =species abundance. Area surveyed in farmland =6.4 km^2^, plantation =5.4 km^2^ and primary forest = 4.8 km^2^. FF=Forest specialists, F=Forest generalists, f=forest visitors. Categorization follows the classification of forest birds by (22).

| Category | Common Name | Scientific Name | Farmland |  |  | Plantation |  |  | Primary forest |  |  |
| --- | --- | --- | --- | --- | --- | --- | --- | --- | --- | --- | --- |
|  |  |  | n | D' | N | n | D' | N | n | D' | N |
| F | Great Sparrowhawk | *Accipiter melanoleucus* | 0 | 0.00 | 0.00 | 2 | 9.44 | 50.96 | 0 | 0.00 | 0.00 |
| F | African Goshawk | *Accipiter tachiro* | 1 | 4.72 | 30.20 | 5 | 23.59 | 127.39 | 2 | 9.44 | 45.29 |
|  | Black-headed Heron | *Ardea melanocephala* | 1 | 4.72 | 30.20 | 0 | 0.00 | 0.00 | 0 | 0.00 | 0.00 |
|  | Hadedda Ibis | *Bostrychia hagedash* | 3 | 14.15 | 90.59 | 0 | 0.00 | 0.00 | 0 | 0.00 | 0.00 |
|  | White-browed Coucal | *Centropus superciliosus* | 15 | 70.77 | 452.94 | 9 | 42.46 | 229.30 | 3 | 14.15 | 67.94 |
| F | Southern Banded Snake-Eagle | *Circaetus fasciolatus* | 0 | 0.00 | 0.00 | 2 | 9.44 | 50.96 | 3 | 14.15 | 67.94 |
| f | African Fish Eagle | *Haliaeetus vocifer* | 4 | 18.87 | 120.78 | 0 | 0.00 | 0.00 | 0 | 0.00 | 0.00 |
| f | Lizard Buzzard | *Kaupifalco monogrammicus* | 16 | 75.49 | 483.13 | 9 | 42.46 | 229.30 | 3 | 14.15 | 67.94 |
|  | Black Kite | *Milvus migrans* | 4 | 18.87 | 120.78 | 3 | 14.15 | 76.43 | 0 | 0.00 | 0.00 |
|  | Sacred Ibis | *Threskiornis aethiopicus* | 1 | 4.72 | 30.20 | 1 | 4.72 | 25.48 | 0 | 0.00 | 0.00 |
| F | Trumpeter Hornbill | *Ceratogymna bucinator* | 12 | 56.62 | 362.35 | 15 | 70.77 | 382.17 | 8 | 37.74 | 181.17 |
| f | Speckled Mousebird | *Colius striatus* | 11 | 51.90 | 332.15 | 2 | 9.44 | 50.96 | 0 | 0.00 | 0.00 |
| F | Fischer's Turaco | *Tauraco fischeri* | 3 | 14.15 | 90.59 | 12 | 56.62 | 305.73 | 3 | 14.15 | 67.94 |
| F | Green Pigeon | *Treron australis* | 16 | 75.49 | 483.13 | 5 | 23.59 | 127.39 | 0 | 0.00 | 0.00 |
|  | Zanzibar Sombre Greenbul | *Andropadus importunus* | 68 | 320.83 | 2053.31 | 30 | 141.54 | 764.33 | 13 | 61.34 | 294.41 |
| F | Yellow-rumped Tinkerbird | *Pogoniulus bilineatus* | 5 | 23.59 | 150.98 | 9 | 42.46 | 229.30 | 4 | 18.87 | 90.59 |
| FF | Eastern Green Tinkerbird | *Pogoniulus simplex* | 0 | 0.00 | 0.00 | 4 | 18.87 | 101.91 | 9 | 42.46 | 203.82 |
| f | Common Bulbul | *Pycnonotus barbatus* | 76 | 358.58 | 2294.88 | 22 | 103.80 | 560.51 | 3 | 14.15 | 67.94 |
| FF | Green Barbet | *Stactolaema olivacea* | 24 | 113.23 | 724.70 | 42 | 198.16 | 1070.06 | 11 | 51.90 | 249.12 |
| f | Crowned Hornbill | *Tockus alboterminatus* | 0 | 0.00 | 0.00 | 2 | 9.44 | 50.96 | 0 | 0.00 | 0.00 |
| FF | Black-bellied Starling | *Notopholia corusca* | 19 | 89.64 | 573.72 | 34 | 160.42 | 866.24 | 38 | 179.29 | 860.58 |
|  | Grey-headed Sparrow | *Passer griseus* | 0 | 0.00 | 0.00 | 3 | 14.15 | 76.43 | 1 | 4.72 | 22.65 |
| f | Yellow-fronted Canary | *Serinus mozambicus* | 35 | 165.13 | 1056.85 | 23 | 108.52 | 585.99 | 0 | 0.00 | 0.00 |
| f | Ring-necked Dove | *Streptopelia capicola* | 14 | 66.05 | 422.74 | 4 | 18.87 | 101.91 | 0 | 0.00 | 0.00 |
| f | Red-eyed Dove | *Streptopelia semitorquata* | 12 | 56.62 | 362.35 | 6 | 28.31 | 152.87 | 0 | 0.00 | 0.00 |
| f | Emerald-spotted Wood-dove | *Turtur chalcospilos* | 21 | 99.08 | 634.11 | 11 | 51.90 | 280.25 | 10 | 47.18 | 226.47 |
| F | Tambourine Dove | *Turtur tympanistria* | 4 | 18.87 | 120.78 | 9 | 42.46 | 229.30 | 8 | 37.74 | 181.17 |
|  | Common Waxbill | *Estrilda astrild* | 0 | 0.00 | 0.00 | 2 | 9.44 | 50.96 | 0 | 0.00 | 0.00 |
|  | Crested Francolin | *Francolinus sephaena* | 1 | 4.72 | 30.20 | 0 | 0.00 | 0.00 | 0 | 0.00 | 0.00 |
| F | Kenya Crested Guineafowl | *Guttera pucherani* | 0 | 0.00 | 0.00 | 0 | 0.00 | 0.00 | 10 | 47.18 | 226.47 |
|  | Red-billed Firefinch | *Lagonosticta senegala* | 1 | 4.72 | 30.20 | 0 | 0.00 | 0.00 | 0 | 0.00 | 0.00 |
|  | Bronze Mannikin | *Lonchura cucullata* | 46 | 217.03 | 1389.01 | 19 | 89.64 | 484.08 | 7 | 33.03 | 158.53 |
|  | Helmeted Guineafowl | *Numida meleagris* | 1 | 4.72 | 30.20 | 0 | 0.00 | 0.00 | 0 | 0.00 | 0.00 |
|  | Black-headed Weaver | *Ploceus cucullatus* | 18 | 84.93 | 543.52 | 2 | 9.44 | 50.96 | 5 | 23.59 | 113.23 |
|  | Red-cheeked Cordon-bleu | *Uraeginthus bengalus* | 12 | 56.62 | 362.35 | 0 | 0.00 | 0.00 | 0 | 0.00 | 0.00 |
| F | Peters's Twinspot | *Hypargos niveoguttatus* | 0 | 0.00 | 0.00 | 9 | 42.46 | 229.30 | 1 | 4.72 | 22.65 |
| FF | Sokoke Pipit | *Anthus sokokensis* | 0 | 0.00 | 0.00 | 0 | 0.00 | 0.00 | 10 | 47.18 | 226.47 |
| FF | Black-headed Apalis | *Apalis melanocephala* | 2 | 9.44 | 60.39 | 9 | 42.46 | 229.30 | 80 | 377.45 | 1811.75 |
| F | Narina Trogon | *Apaloderma narina* | 0 | 0.00 | 0.00 | 3 | 14.15 | 76.43 | 6 | 28.31 | 135.88 |
| FF | Forest Batis | *Batis mixta* | 2 | 9.44 | 60.39 | 0 | 0.00 | 0.00 | 17 | 80.21 | 385.00 |
| F | Pale Batis | *Batis soror* | 0 | 0.00 | 0.00 | 0 | 0.00 | 0.00 | 25 | 117.95 | 566.17 |
|  | Green-backed Camaroptera | *Camaroptera brachyura* | 0 | 0.00 | 0.00 | 1 | 4.72 | 25.48 | 1 | 4.72 | 22.65 |
| f | Grey-backed Camaroptera | *Camaroptera brevicaudata* | 12 | 56.62 | 362.35 | 21 | 99.08 | 535.03 | 36 | 169.85 | 815.29 |
| f | Black Cuckoo-shrike | *Campephaga flava* | 1 | 4.72 | 30.20 | 3 | 14.15 | 76.43 | 4 | 18.87 | 90.59 |
| F | Mombasa Woodpecker | *Campethera mombassica* | 3 | 14.15 | 90.59 | 10 | 47.18 | 254.78 | 9 | 42.46 | 203.82 |
| f | Eastern Bearded Scrub-Robin | *Cercotrichas quadrivirgata* | 0 | 0.00 | 0.00 | 5 | 23.59 | 127.39 | 14 | 66.05 | 317.06 |
| F | Yellowbill | *Ceuthmochares aereus* | 4 | 18.87 | 120.78 | 9 | 42.46 | 229.30 | 2 | 9.44 | 45.29 |
| f | Klaas's Cuckoo | *Chrysococcyx klaas* | 0 | 0.00 | 0.00 | 0 | 0.00 | 0.00 | 1 | 4.72 | 22.65 |
| f | White-browed Robin-Chat | *Cossypha heuglini* | 7 | 33.03 | 211.37 | 18 | 84.93 | 458.60 | 9 | 42.46 | 203.82 |
| F | Red-capped Robin-Chat | *Cossypha natalensis* | 8 | 37.74 | 241.57 | 23 | 108.52 | 585.99 | 16 | 75.49 | 362.35 |
|  | African Palm Swift | *Cypsiurus parvus* | 5 | 23.59 | 150.98 | 2 | 9.44 | 50.96 | 6 | 28.31 | 135.88 |
|  | Fork-tailed Drongo | *Dicrurus adsimilis* | 44 | 207.60 | 1328.62 | 23 | 108.52 | 585.99 | 12 | 56.62 | 271.76 |
| F | Black-backed Puffback | *Dryocopus cubla* | 23 | 108.52 | 694.50 | 8 | 37.74 | 203.82 | 19 | 89.64 | 430.29 |
| FF | Little Yellow Flycatcher | *Erythrocerus holochlorus* | 0 | 0.00 | 0.00 | 22 | 103.80 | 560.51 | 34 | 160.42 | 769.99 |
| f | Grey-headed Kingfisher | *Halcyon leucocephala* | 1 | 4.72 | 30.20 | 0 | 0.00 | 0.00 | 0 | 0.00 | 0.00 |
|  | Mangrove Kingfisher | *Halcyon senegaloides* | 2 | 9.44 | 60.39 | 3 | 14.15 | 76.43 | 2 | 9.44 | 45.29 |
|  | Lesser Striped Swallow | *Hirundo abyssinica* | 0 | 0.00 | 0.00 | 6 | 28.31 | 152.87 | 0 | 0.00 | 0.00 |
| f | Pallid Honeyguide | *Indicator meliphilus* | 0 | 0.00 | 0.00 | 11 | 51.90 | 280.25 | 1 | 4.72 | 22.65 |
| f | Lesser Honeyguide | *Indicator minor* | 0 | 0.00 | 0.00 | 6 | 28.31 | 152.87 | 0 | 0.00 | 0.00 |
| f | Scaly-throated Honeyguide | *Indicator variegatus* | 0 | 0.00 | 0.00 | 8 | 37.74 | 203.82 | 2 | 9.44 | 45.29 |
| f | African Pygmy Kingfisher | *Ispidina lecontei* | 0 | 0.00 | 0.00 | 0 | 0.00 | 0.00 | 1 | 4.72 | 22.65 |
| f | Tropical Boubou | *Laniarius aethiopicus* | 30 | 141.54 | 905.87 | 13 | 61.34 | 331.21 | 35 | 165.13 | 792.64 |
|  | Slate-coloured Boubou | *Laniarius funebris* | 0 | 0.00 | 0.00 | 0 | 0.00 | 0.00 | 1 | 4.72 | 22.65 |
| F | Four-coloured Bush-shrike | *Malaconotus quadricolor* | 0 | 0.00 | 0.00 | 0 | 0.00 | 0.00 | 21 | 99.08 | 475.58 |
| f | White-throated Bee-eater | *Merops albicollis* | 0 | 0.00 | 0.00 | 5 | 23.59 | 127.39 | 2 | 9.44 | 45.29 |
|  | Northern Carmine Bee-eater | *Merops nubicus* | 2 | 9.44 | 60.39 | 0 | 0.00 | 0.00 | 0 | 0.00 | 0.00 |
|  | African Pied Wagtail | *Motacilla aguimp* | 8 | 37.74 | 241.57 | 1 | 4.72 | 25.48 | 0 | 0.00 | 0.00 |
| F | Ashy Flycatcher | *Muscicapa caerulea* | 2 | 9.44 | 60.39 | 0 | 0.00 | 0.00 | 2 | 9.44 | 45.29 |
| FF | Red-tailed Ant-Thrush | *Neocossyphus rufus* | 0 | 0.00 | 0.00 | 4 | 18.87 | 101.91 | 13 | 61.34 | 294.41 |
|  | Eastern Nicator | *Nicator gularis* | 3 | 14.15 | 90.59 | 11 | 51.90 | 280.25 | 22 | 103.80 | 498.23 |
| f | African Golden Oriole | *Oriolus auratus* | 10 | 47.18 | 301.96 | 15 | 70.77 | 382.17 | 0 | 0.00 | 0.00 |
| f | Black-headed Oriole | *Oriolus larvatus* | 12 | 56.62 | 362.35 | 23 | 108.52 | 585.99 | 12 | 56.62 | 271.76 |
| f | Eurasian Golden Oriole | *Oriolus oriolus* | 1 | 4.72 | 30.20 | 0 | 0.00 | 0.00 | 0 | 0.00 | 0.00 |
| f | Thick-billed Cuckoo | *Pachycoccyx audeberti* | 27 | 127.39 | 815.29 | 0 | 0.00 | 0.00 | 0 | 0.00 | 0.00 |
|  | Green Wood-hoopoe | *Phoeniculus purpureus* | 0 | 0.00 | 0.00 | 3 | 14.15 | 76.43 | 1 | 4.72 | 22.65 |
| FF | Tiny Greenbul | *Phyllastrephus debilis* | 0 | 0.00 | 0.00 | 0 | 0.00 | 0.00 | 23 | 108.52 | 520.88 |
| FF | Fischer's Greenbul | *Phyllastrephus fischeri* | 0 | 0.00 | 0.00 | 2 | 9.44 | 50.96 | 54 | 254.78 | 1222.93 |
| f | Northern Brownbul | *Phyllastrephus strepitans* | 6 | 28.31 | 181.17 | 28 | 132.11 | 713.38 | 12 | 56.62 | 271.76 |
| F | Terrestrial Brownbul | *Phyllastrephus terrestris* | 0 | 0.00 | 0.00 | 5 | 23.59 | 127.39 | 13 | 61.34 | 294.41 |
| F | Dark-backed Weaver | *Ploceus bicolor* | 2 | 9.44 | 60.39 | 8 | 37.74 | 203.82 | 23 | 108.52 | 520.88 |
| FF | Clarke's Weaver | *Ploceus golandi* | 0 | 0.00 | 0.00 | 0 | 0.00 | 0.00 | 4 | 18.87 | 90.59 |
| f | Tawny-flanked Prinia | *Prinia subflava* | 51 | 240.62 | 1539.99 | 9 | 42.46 | 229.30 | 0 | 0.00 | 0.00 |
| f | Retz's Helmet-shrike | *Prionops retzii* | 0 | 0.00 | 0.00 | 15 | 70.77 | 382.17 | 4 | 18.87 | 90.59 |
| F | Chestnut-fronted Helmet-shrike | *Prionops scopifrons* | 0 | 0.00 | 0.00 | 19 | 89.64 | 484.08 | 75 | 353.86 | 1698.51 |
|  | Common Scimitarbill | *Rhinopomastus cyanomelas* | 1 | 4.72 | 30.20 | 4 | 18.87 | 101.91 | 5 | 23.59 | 113.23 |
| FF | East Coast Akalat | *Sheppardia gunningi* | 0 | 0.00 | 0.00 | 0 | 0.00 | 0.00 | 11 | 51.90 | 249.12 |
|  | Black-crowned Tchagra | *Tchagra australis* | 18 | 84.93 | 543.52 | 2 | 9.44 | 50.96 | 5 | 23.59 | 113.23 |
| f | African Paradise-flycatcher | *Terpsiphone viridis* | 3 | 14.15 | 90.59 | 7 | 33.03 | 178.34 | 3 | 14.15 | 67.94 |
| FF | Blue-mantled Crested-flycatcher | *Trochocercus cyanomelas* | 0 | 0.00 | 0.00 | 0 | 0.00 | 0.00 | 17 | 80.21 | 385.00 |
|  | Scaly Babbler | *Turdoides squamulatus* | 31 | 146.26 | 936.07 | 48 | 226.47 | 1222.93 | 0 | 0.00 | 0.00 |
| FF | Amani Sunbird | *Anthreptes pallidigaster* | 0 | 0.00 | 0.00 | 4 | 18.87 | 101.91 | 22 | 103.80 | 498.23 |
| FF | Plain-backed Sunbird | *Anthreptes reichenowi* | 0 | 0.00 | 0.00 | 1 | 4.72 | 25.48 | 10 | 47.18 | 226.47 |
| F | Collared Sunbird | *Hedydipna collaris* | 41 | 193.44 | 1238.03 | 53 | 250.06 | 1350.32 | 61 | 287.80 | 1381.46 |
| FF | Olive Sunbird | *Nectarinia olivacea* | 21 | 99.08 | 634.11 | 32 | 150.98 | 815.29 | 61 | 287.80 | 1381.46 |
| f | Amethyst Sunbird | *Nectarinia amethystina* | 1 | 4.72 | 30.20 | 0 | 0.00 | 0.00 | 0 | 0.00 | 0.00 |
|  | Pied Crow | *Corvus albus* | 8 | 37.74 | 241.57 | 0 | 0.00 | 0.00 | 0 | 0.00 | 0.00 |
|  | House Crow | *Corvus splendens* | 7 | 33.03 | 211.37 | 0 | 0.00 | 0.00 | 0 | 0.00 | 0.00 |

### Avian community similarity

The primary forest holds a different bird community compared to plantation forest and farmland. Only 11 sampling points showed similarity in species composition between plantation forest and primary forest despite their close proximity, whereas 44 points showed similarity between plantations and farmland. Primary forest and farmland shared few species other than Collared Sunbird and Zanzibar Sombre Greenbul that were well distributed across all three land use types.

### Effect of habitat factors

Vertical vegetation heterogeneity had a positive influence on overall bird species richness. However, the number of large trees had no significant influence (ns), R^2^= 0.039231. The best habitat model for increased richness of bird species in farmlands was obtained from a mixed effect model with the number of trees with fruits and the number of large trees based on Akaike’s Information Criterion (AIC). However, the differences in AIC values were small, an indication that all the models could be applied with possible similar effects (Table 2).

**Table 2:** Akaike’s Information Criterion (AIC) of different models for the mixed effect of habitat factors. Many trees and fruiting trees were counts of individuals. Vertical vegetation heterogeneity was based on the Shannon diversity index of % cover of vegetation layers. The models are ranked based on the AIC value. The lowest AIC value indicates the best model.

| Mixed effect model of habitat factors | AIC value | Deviance |
| --- | --- | --- |
| Farmland + Fruit trees + Large trees | 159.58 | 28.401 |
| Farmland + Fruiting trees + Large trees + Vertical vegetation heterogeneity | 161.1 | 28.173 |
| Farmland + Fruiting trees + Large trees + Proximity to settlement | 161.22 | 28.272 |
| Farmland + Fruiting trees + Large trees + Proximity to forest | 161.58 | 28.404 |
| Farmland + Fruiting trees + vertical vegetation heterogeneity | 162.89 | 29.011 |

## DISCUSSION

### Response of the avian community to land use changes

As expected, bird species diversity was highest in the primary forest while the farmlands supported fewer species. This has been reported in many other surveys, e.g. (9,30,31). High bird species diversity in the primary forest can be attributed in part to the complexity of habitat structure that provided resources for feeding and nesting (32), high vertical vegetation heterogeneity and the presence of fruiting trees. The primary forest at Arabuko Sokoke consisted of diverse vegetation layers ranging from the understorey to the upper canopy. With similar sampling effort in the three vegetation types in the primary forest, we recorded greater species diversity in *Brachystegia* and Mixed Forest, and lower diversity in the *Cynometra* zone. This could be partly explained by the difference in vertical vegetation heterogeneity observed between the vegetation types. *Brachystegia* and Mixed Forest had many large trees and a dense understory, whereas the *Cynometra* thicket had no large trees and vegetation at a uniform height. However, there was limited species overlap between the bird community in the primary forest and the nearby plantation forest. Whereas points in the primary forest were characterized by high vertical vegetation heterogeneity and a high number of fruiting trees, the plantation forest was characterized by wide open spaces, low tree density and a low number of fruiting trees. Similarly, (33) in tropical montane cloud forest in Guatemala; (34) in indigenous forest reserves of Talamanca, Costa Rica, and (35) in West Java, Indonesia noted a marked reduction in forest species and an overall shift in bird species composition in agroforestry systems. Some forest specialists (FF) bird species in Arabuko Sokoke appear to be highly specialized and sparsely distributed e.g. Pale Batis, Tiny Greenbul, Four-coloured Bush-shrike; and are wholly dependent on the forest for nesting, food resources and refuge from predators. Forest interior species have been reported to be highly sensitive to disturbance and will even shun clearings or gaps resulting from treefalls (36) logged areas may take many years to be recolonised (37) and species breeding in holes or crevices are particularly adversely affected (38).

The decline in bird species diversity in plantation forest and farmlands can be linked to the extent of disturbance and reduction of native vegetation in these two land use types. In farmlands around the forest, there are few native fruiting plant species on which birds and other animals may depend, so that farmlands support fewer species (9,39). Low canopy cover in plantations and farmlands due to intensive disturbance leading to low vertical vegetation heterogeneity, could be a major contributor to the low number of species and the reduced density and abundance of many bird species. However, we found evidence of increased diversity at points in farmlands with remnant native vegetation patches. Other studies have also shown that even small changes in the structure and composition of tree cover may have a significant impact on bird assemblages (5,32), leading to changes in bird diversity and community composition, with fewer species and foraging guilds present in more intensively managed landscapes (7). Logging for timber in the plantation forest resulting in clear-felled areas could also be a contributing factor to low species counts. Excessive disturbance brought about by logging operations usually reduces total avian diversity (40) or promotes an influx of non-forest species which replace forest specialists (37). Such changes particularly affect less abundant, range-restricted birds and rainforest specialists (8). The trees in the plantations, mainly *Eucalyptus sp*. and *Casuarina sp*. provide few food resources. In East Usambara, Tanzania, (41) also recorded a significantly lower diversity of forest birds in *Eucalyptus* plantations compared to primary forest, which they attributed to limited nesting opportunities and reduced understorey cover. Plantation forest in this study was associated with reduced variation in the vegetation structure, which lacks an intermediate stratum which many bird species require.

Planting of native tree species is unlikely to be adopted by farmers without either financial incentives, or clear evidence that they provide additional benefits (42). Farmers plant a few exotic fruiting trees like the neem tree, cashew nuts and mangoes for their household use. These are then also utilized by frugivores, but there are no incentives to retain native vegetation in farmlands. Native forest patches create habitat heterogeneity and provide foraging and roosting sites (43), resources that are scarce in the open farmlands (44).

The General Linear Modelling analysis of this study found a significant positive influence of the mixed effect of the number of large trees and the number of fruiting trees on bird diversity. Heterogeneous vegetation structure will provide a range of perching heights for insectivorous bird species including forest specialists (FF) like Plain-backed Sunbird, Sokoke Pipit, Pale Batis, Blue-mantled Crested-flycatcher, and Clarke’s Weaver. The importance of understorey vegetation for bird diversity was also emphasized by (45) in the Kakamega forest in western Kenya. Thus, the structure of the habitat or land use system may be more important to many bird species than the plant species composition. The importance of habitat structure for forest birds has been shown elsewhere, e.g. (5) in Talamanca, Costa Rica, (46) on land-bridge forest islands in Peninsular, Malaysia, (45) at the Kakamega forest in Western Kenya, and (47) on Sitka spruce plantations in Ireland.

While conservation of the primary forest is important for bird conservation in this area, breeding areas outside the forest may be equally significant. The breeding of Clarke’s Weaver had been a mystery for many years and the nest had never been described (48). Only recently the birds were discovered nesting low down within the sedges of a seasonal wetland in the Dakatcha woodland some 60 km from the forest (49). While the Dakatcha woodland is a designated Important Bird and Biodiversity Area (IBA), it has no formal protection, but falls under the management of a local community conservation group, and the site is used for harvesting thatching material (49). Formal protection of this woodland and the adjacent wetland is thus essential for preserving the breeding sites of this endemic species, which at other times is apparently restricted to Arabuko Sokoke Forest.

### Response of birds to habitat factors

Some species of special conservation concern were forest restricted including the Sokoke Pipit, Clarke’s Weaver and the Kenya Crested Guineafowl (*Guttera pucherani*) while others showed strong forest dependence (Appendix 1). Similarly in the Solomon Islands endemic species were largely restricted to the forest, whereas birds with a wider distribution were more common in other land-use types (50). Farmland and plantations around Arabuko Sokoke Forest are currently subject to frequent large scale disturbance and a decline of forest specialists in such degraded farmland has also been reported by (51). Forest specialists species are typically the first to be lost from degraded areas, whereas widespread, generalist species may increase in abundance (52). The decrease in abundance of forest specialists in plantation forest despite the closeness to the primary forest could be indicative of the unsuitability of *Eucalyptus* sp. which dominates the plantation stands to use by birds. This agrees with studies in other regions that documented that forest specialists diminished in abundance in plantations e.g. (53). However, the occurrence some Forest specialist species (FF) in large numbers across the three habitats, confirms the usefulness of farmlands in bird conservation, as in the case of Budongo forest in Uganda where (22) reported some bird species to have more extensive distributions in the neighbouring farmlands. Our findings focus for the first time on a combination of land use variables that will promote bird utilization of farmlands around ASF. Native vegetation cover in farmland has been greatly reduced for small scale subsistence agriculture with only a few tree stands and narrow strips of remnant natural vegetation. Promoting planting of short duration nitrogen fixing trees where crops are planted could motivate farmers to adopt on-farm planting of trees. Such trees improve soil fertility and are useful in the short term as fuel wood, hence providing multiple benefits to the farmers (54). Farmers can also be influenced to adopt on-farm tree planting by supplying them with quality tree seeds and involving them in experimental agroforestry alongside researchers (55). Redesigning agricultural landscapes to be more diverse could promote both crop production, ecosystem services such as pollination (56–58) and use by forest birds. Resident forest species are often reluctant to cross open farmlands, which thus act as barriers for their dispersal (59). A marked decrease in forest species composition along a gradient from primary forest to farmland has also been reported in South West Cameroon (1) and in Sumatra, where up to half of the total number of forest birds was lost in open farmlands (35).

## Conclusions

We found that Arabuko Sokoke Forest has a bird community distinct from both the neighbouring plantation forest and farmlands. Patterns of habitat use by birds in the area suggest that vertical vegetation heterogeneity is especially significant in sustaining diverse and abundant bird populations, while pure stand plantations with high levels of disturbance attract fewer bird species even if they are near native forests. Improving the vegetation structure by retaining both large emergent trees and an understorey could increase the habitat quality of plantations and farmlands for many birds. Tree density in plantation forest could be increased by planting high value and fast-growing trees and limiting clear-fell harvesting methods that lead to large open gaps, which would improve connectivity with primary forest, while habitat quality in farmlands could be improved by maintaining stands of large trees and increasing the number of fruiting trees. For an integrated conservation plan, we urgently need information on local movement patterns of bird populations, and the current extent of connectivity between the remaining patches of coastal forest.

## ACKNOWLEDGEMENTS

The Kenya Forest Service granted permission for the research in Arabuko Sokoke Forest, and the local community allowed the surveys on their farms. Many thanks to the International Foundation for Science (IFS) for funding the research and Nature Kenya for administering the funds. Rhodes University provided off-campus access to library resources and a platform for research and Strathmore University essential logistical support. Rose Njuguna and Felex Namayi assisted in updating categories in the species list.

